## Supplemental file for "The intrinsically disordered cytoplasmic tail of a dendrite branching receptor uses two distinct mechanisms to regulate the actin cytoskeleton"

**Supplementary File 1: Amino acid sequences of recombinant proteins used in this study.**

| Construct name | Description | Plasmid Identity | Sequence | Source or reference |
| --- | --- | --- | --- | --- |
| GST-ICD | GST-thrombin-TEV-ceHPO-30 ICD (a.a. 229-279) | pDK079 | MSPILGYWKIKGLVQPTRLLLEYLEEKYEEHLYERDEGDKWRNKKFELGLEFPNLPYYIDGDVKLTQSMAIIRYIADKHNMLGGCPKERAEISMLEGAVLDIRYGVSRIAYSKDFETLKVDFLSKLPEMLKMFEDRLCHKTYLNGDHVTHPDFMLYDALDVVLYMDPMCLDAFPKLVCFKKRIEAIPQIDKYLKSSKYIAWPLQGWQATFGGGDHPPKSDLVPRGSENLYFQGHM**TSKHAHDVCCTSRKEYREQTKWKNNGLILKTGRVNHQSHRPFVVIDDDSSM** | Zou, 2018 |
| GST-ICD ∆1 | GST- thrombin-TEV-ceHPO-30 ICD Ala 229-233 | pDK325 | MSPILGYWKIKGLVQPTRLLLEYLEEKYEEHLYERDEGDKWRNKKFELGLEFPNLPYYIDGDVKLTQSMAIIRYIADKHNMLGGCPKERAEISMLEGAVLDIRYGVSRIAYSKDFETLKVDFLSKLPEMLKMFEDRLCHKTYLNGDHVTHPDFMLYDALDVVLYMDPMCLDAFPKLVCFKKRIEAIPQIDKYLKSSKYIAWPLQGWQATFGGGDHPPKSDLVPRGSENLYFQGHM**AAAAAHDVCCTSRKEYREQTKWKNNGLILKTGRVNHQSHRPFVVIDDDSSM** | Zou, 2018 |
| GST-ICD ∆2 | GST- thrombin-TEV-ceHPO-30 ICD Ala 234-238 | pDK326 | MSPILGYWKIKGLVQPTRLLLEYLEEKYEEHLYERDEGDKWRNKKFELGLEFPNLPYYIDGDVKLTQSMAIIRYIADKHNMLGGCPKERAEISMLEGAVLDIRYGVSRIAYSKDFETLKVDFLSKLPEMLKMFEDRLCHKTYLNGDHVTHPDFMLYDALDVVLYMDPMCLDAFPKLVCFKKRIEAIPQIDKYLKSSKYIAWPLQGWQATFGGGDHPPKSDLVPRGSENLYFQGHM**TSKHAAAAAATSRKEYREQTKWKNNGLILKTGRVNHQSHRPFVVIDDDSSM** | Zou, 2018 |
| GST-ICD ∆3 | GST- thrombin-TEV-ceHPO-30 ICD Ala 239-243 | pDK327 | MSPILGYWKIKGLVQPTRLLLEYLEEKYEEHLYERDEGDKWRNKKFELGLEFPNLPYYIDGDVKLTQSMAIIRYIADKHNMLGGCPKERAEISMLEGAVLDIRYGVSRIAYSKDFETLKVDFLSKLPEMLKMFEDRLCHKTYLNGDHVTHPDFMLYDALDVVLYMDPMCLDAFPKLVCFKKRIEAIPQIDKYLKSSKYIAWPLQGWQATFGGGDHPPKSDLVPRGSENLYFQGHM**TSKHAHDVCCAAAAAYREQTKWKNNGLILKTGRVNHQSHRPFVVIDDDSSM** | Zou, 2018 |
| GST-ICD ∆4 | GST- thrombin-TEV-ceHPO-30 ICD Ala 244-248 | pDK328 | MSPILGYWKIKGLVQPTRLLLEYLEEKYEEHLYERDEGDKWRNKKFELGLEFPNLPYYIDGDVKLTQSMAIIRYIADKHNMLGGCPKERAEISMLEGAVLDIRYGVSRIAYSKDFETLKVDFLSKLPEMLKMFEDRLCHKTYLNGDHVTHPDFMLYDALDVVLYMDPMCLDAFPKLVCFKKRIEAIPQIDKYLKSSKYIAWPLQGWQATFGGGDHPPKSDLVPRGSENLYFQGHM**TSKHAHDVCCTSRKEAAAAAKWKNNGLILKTGRVNHQSHRPFVVIDDDSSM** | Zou, 2018 |
| GST-ICD ∆5 | GST- thrombin-TEV-ceHPO-30 ICD Ala 249-253 | pDK329 | MSPILGYWKIKGLVQPTRLLLEYLEEKYEEHLYERDEGDKWRNKKFELGLEFPNLPYYIDGDVKLTQSMAIIRYIADKHNMLGGCPKERAEISMLEGAVLDIRYGVSRIAYSKDFETLKVDFLSKLPEMLKMFEDRLCHKTYLNGDHVTHPDFMLYDALDVVLYMDPMCLDAFPKLVCFKKRIEAIPQIDKYLKSSKYIAWPLQGWQATFGGGDHPPKSDLVPRGSENLYFQGHM**TSKHAHDVCCTSRKEYREQTAAAAAGLILKTGRVNHQSHRPFVVIDDDSSM** | Zou, 2018 |
| GST-ICD ∆6 | GST- thrombin-TEV-ceHPO-30 ICD Ala 254-258 | pDK330 | MSPILGYWKIKGLVQPTRLLLEYLEEKYEEHLYERDEGDKWRNKKFELGLEFPNLPYYIDGDVKLTQSMAIIRYIADKHNMLGGCPKERAEISMLEGAVLDIRYGVSRIAYSKDFETLKVDFLSKLPEMLKMFEDRLCHKTYLNGDHVTHPDFMLYDALDVVLYMDPMCLDAFPKLVCFKKRIEAIPQIDKYLKSSKYIAWPLQGWQATFGGGDHPPKSDLVPRGSENLYFQGHM**TSKHAHDVCCTSRKEYREQTKWKNNAAAAATGRVNHQSHRPFVVIDDDSSM** | Zou, 2018 |
| GST-ICD ∆7 | GST- thrombin-TEV-ceHPO-30 ICD Ala 259-263 | pDK331 | MSPILGYWKIKGLVQPTRLLLEYLEEKYEEHLYERDEGDKWRNKKFELGLEFPNLPYYIDGDVKLTQSMAIIRYIADKHNMLGGCPKERAEISMLEGAVLDIRYGVSRIAYSKDFETLKVDFLSKLPEMLKMFEDRLCHKTYLNGDHVTHPDFMLYDALDVVLYMDPMCLDAFPKLVCFKKRIEAIPQIDKYLKSSKYIAWPLQGWQATFGGGDHPPKSDLVPRGSENLYFQGHM**TSKHAHDVCCTSRKEYREQTKWKNNGLILKAAAAAHQSHRPFVVIDDDSSM** | Zou, 2018 |
| ICD-GST | ceHPO-30 ICD (a.a. 229-279)-GG-3C-GST | pDK238 | M**TSKHAHDVCCTSRKEYREQTKWKNNGLILKTGRVNHQSHRPFVVIDDDSSM**GGLEVLFQGPMSPILGYWKIKGLVQPTRLLLEYLEEKYEEHLYERDEGDKWRNKKFELGLEFPNLPYYIDGDVKLTQSMAIIRYIADKHNMLGGCPKERAEISMLEGAVLDIRYGVSRIAYSKDFETLKVDFLSKLPEMLKMFEDRLCHKTYLNGDHVTHPDFMLYDALDVVLYMDPMCLDAFPKLVCFKKRIEAIPQIDKYLKSSKYIAWPLQGWQATFGGGDHPPK | This paper |
| ICD ∆8 – GST | ceHPO-30 ICD Ala 264-268 | pDK257 | M**TSKHAHDVCCTSRKEYREQTKWKNNGLILKTGRVNAAAAAPFVVIDDDSSM**GGLEVLFQGPMSPILGYWKIKGLVQPTRLLLEYLEEKYEEHLYERDEGDKWRNKKFELGLEFPNLPYYIDGDVKLTQSMAIIRYIADKHNMLGGCPKERAEISMLEGAVLDIRYGVSRIAYSKDFETLKVDFLSKLPEMLKMFEDRLCHKTYLNGDHVTHPDFMLYDALDVVLYMDPMCLDAFPKLVCFKKRIEAIPQIDKYLKSSKYIAWPLQGWQATFGGGDHPPK | This paper |
| ICD ∆9 - GST | ceHPO-30 ICD Ala 269-273 | pDK258 | M**TSKHAHDVCCTSRKEYREQTKWKNNGLILKTGRVNHQSHRAAAAADDDSSM**GGLEVLFQGPMSPILGYWKIKGLVQPTRLLLEYLEEKYEEHLYERDEGDKWRNKKFELGLEFPNLPYYIDGDVKLTQSMAIIRYIADKHNMLGGCPKERAEISMLEGAVLDIRYGVSRIAYSKDFETLKVDFLSKLPEMLKMFEDRLCHKTYLNGDHVTHPDFMLYDALDVVLYMDPMCLDAFPKLVCFKKRIEAIPQIDKYLKSSKYIAWPLQGWQATFGGGDHPPK | This paper |
| ICD ∆10 - GST | ceHPO-30 ICD Ala 274-279 | pDK259 | M**TSKHAHDVCCTSRKEYREQTKWKNNGLILKTGRVNHQSHRPFVVIAAAAAA**GGLEVLFQGPMSPILGYWKIKGLVQPTRLLLEYLEEKYEEHLYERDEGDKWRNKKFELGLEFPNLPYYIDGDVKLTQSMAIIRYIADKHNMLGGCPKERAEISMLEGAVLDIRYGVSRIAYSKDFETLKVDFLSKLPEMLKMFEDRLCHKTYLNGDHVTHPDFMLYDALDVVLYMDPMCLDAFPKLVCFKKRIEAIPQIDKYLKSSKYIAWPLQGWQATFGGGDHPPK | This paper |
| GST-ICD ∆11 | GST- thrombin-TEV-ceHPO-30 ICD GGS 249-258 | pDK332 | MSPILGYWKIKGLVQPTRLLLEYLEEKYEEHLYERDEGDKWRNKKFELGLEFPNLPYYIDGDVKLTQSMAIIRYIADKHNMLGGCPKERAEISMLEGAVLDIRYGVSRIAYSKDFETLKVDFLSKLPEMLKMFEDRLCHKTYLNGDHVTHPDFMLYDALDVVLYMDPMCLDAFPKLVCFKKRIEAIPQIDKYLKSSKYIAWPLQGWQATFGGGDHPPKSDLVPRGSENLYFQGHM**TSKHAHDVCCTSRKEYREQTGGSGGSGGSGTGRVNHQSHRPFVVIDDDSSM** | Zou, 2018 |
| GST-ICD ∆12 | GST- thrombin-TEV-ceHPO-30 ICD GGS 264-273 | pDK333 | MSPILGYWKIKGLVQPTRLLLEYLEEKYEEHLYERDEGDKWRNKKFELGLEFPNLPYYIDGDVKLTQSMAIIRYIADKHNMLGGCPKERAEISMLEGAVLDIRYGVSRIAYSKDFETLKVDFLSKLPEMLKMFEDRLCHKTYLNGDHVTHPDFMLYDALDVVLYMDPMCLDAFPKLVCFKKRIEAIPQIDKYLKSSKYIAWPLQGWQATFGGGDHPPKSDLVPRGSENLYFQGHM**TSKHAHDVCCTSRKEYREQTKWKNNGLILKTGRVNGGSGGSGGSGDDDSSM** | Zou, 2018 |
| GST-ICD ∆13 | GST- thrombin-TEV-ceHPO-30 ICD GGS 264-275 | pDK334 | MSPILGYWKIKGLVQPTRLLLEYLEEKYEEHLYERDEGDKWRNKKFELGLEFPNLPYYIDGDVKLTQSMAIIRYIADKHNMLGGCPKERAEISMLEGAVLDIRYGVSRIAYSKDFETLKVDFLSKLPEMLKMFEDRLCHKTYLNGDHVTHPDFMLYDALDVVLYMDPMCLDAFPKLVCFKKRIEAIPQIDKYLKSSKYIAWPLQGWQATFGGGDHPPKSDLVPRGSENLYFQGHM**TSKHAHDVCCTSRKEYREQTGGSGGSGGSGGSGGSGGSGGSGGSGGSDSSM** | Zou, 2018 |
| GST-ICD ∆14 | GST- thrombin-TEV-ceHPO-30 ICD GGS 229-248 | pDK335 | MSPILGYWKIKGLVQPTRLLLEYLEEKYEEHLYERDEGDKWRNKKFELGLEFPNLPYYIDGDVKLTQSMAIIRYIADKHNMLGGCPKERAEISMLEGAVLDIRYGVSRIAYSKDFETLKVDFLSKLPEMLKMFEDRLCHKTYLNGDHVTHPDFMLYDALDVVLYMDPMCLDAFPKLVCFKKRIEAIPQIDKYLKSSKYIAWPLQGWQATFGGGDHPPKSDLVPRGSENLYFQGHM**GGSGGSGGSGGSGGSGGSGGKWKNNGLILKTGRVNHQSHRPFVVIDDDSSM** | Zou, 2018 |
| GST-ICD black | GST-thrombin-TEV-ceHPO-30 ICD Y245I F271L | pDK356 | MSPILGYWKIKGLVQPTRLLLEYLEEKYEEHLYERDEGDKWRNKKFELGLEFPNLPYYIDGDVKLTQSMAIIRYIADKHNMLGGCPKERAEISMLEGAVLDIRYGVSRIAYSKDFETLKVDFLSKLPEMLKMFEDRLCHKTYLNGDHVTHPDFMLYDALDVVLYMDPMCLDAFPKLVCFKKRIEAIPQIDKYLKSSKYIAWPLQGWQATFGGGDHPPKSDLVPRGSENLYFQGHM**TSKHAHDVCCTSRKEIREQTKWKNNGLILKTGRVNHQSHRPLVVIDDDSSM** | This paper |
| GST-ICD black W22L | GST-thrombin-TEV-ceHPO-30 ICD Y245I W251L F271L | pDK357 | MSPILGYWKIKGLVQPTRLLLEYLEEKYEEHLYERDEGDKWRNKKFELGLEFPNLPYYIDGDVKLTQSMAIIRYIADKHNMLGGCPKERAEISMLEGAVLDIRYGVSRIAYSKDFETLKVDFLSKLPEMLKMFEDRLCHKTYLNGDHVTHPDFMLYDALDVVLYMDPMCLDAFPKLVCFKKRIEAIPQIDKYLKSSKYIAWPLQGWQATFGGGDHPPKSDLVPRGSENLYFQGHM**TSKHAHDVCCTSRKEIREQTKLKNNGLILKTGRVNHQSHRPLVVIDDDSSM** | This paper |
| GST-ICD black W22M | GST-thrombin-TEV-ceHPO-30 ICD Y245I W251M F271L | pDK358 | MSPILGYWKIKGLVQPTRLLLEYLEEKYEEHLYERDEGDKWRNKKFELGLEFPNLPYYIDGDVKLTQSMAIIRYIADKHNMLGGCPKERAEISMLEGAVLDIRYGVSRIAYSKDFETLKVDFLSKLPEMLKMFEDRLCHKTYLNGDHVTHPDFMLYDALDVVLYMDPMCLDAFPKLVCFKKRIEAIPQIDKYLKSSKYIAWPLQGWQATFGGGDHPPKSDLVPRGSENLYFQGHM**TSKHAHDVCCTSRKEIREQTKMKNNGLILKTGRVNHQSHRPLVVIDDDSSM** | This paper |
| GST-ICD black W22I | GST-thrombin-TEV-ceHPO-30 ICD Y245I W251I F271L | pDK359 | MSPILGYWKIKGLVQPTRLLLEYLEEKYEEHLYERDEGDKWRNKKFELGLEFPNLPYYIDGDVKLTQSMAIIRYIADKHNMLGGCPKERAEISMLEGAVLDIRYGVSRIAYSKDFETLKVDFLSKLPEMLKMFEDRLCHKTYLNGDHVTHPDFMLYDALDVVLYMDPMCLDAFPKLVCFKKRIEAIPQIDKYLKSSKYIAWPLQGWQATFGGGDHPPKSDLVPRGSENLYFQGHM**TSKHAHDVCCTSRKEIREQTKIKNNGLILKTGRVNHQSHRPLVVIDDDSSM** | This paper |
| GST-ICD black W22P | GST-thrombin-TEV-ceHPO-30 ICD Y245I W251P F271L | pDK360 | MSPILGYWKIKGLVQPTRLLLEYLEEKYEEHLYERDEGDKWRNKKFELGLEFPNLPYYIDGDVKLTQSMAIIRYIADKHNMLGGCPKERAEISMLEGAVLDIRYGVSRIAYSKDFETLKVDFLSKLPEMLKMFEDRLCHKTYLNGDHVTHPDFMLYDALDVVLYMDPMCLDAFPKLVCFKKRIEAIPQIDKYLKSSKYIAWPLQGWQATFGGGDHPPKSDLVPRGSENLYFQGHM**TSKHAHDVCCTSRKEIREQTKPKNNGLILKTGRVNHQSHRPLVVIDDDSSM** | This paper |
| GST-ICD black W22H | GST-thrombin-TEV-ceHPO-30 ICD Y245I W251H F271L | pDK361 | MSPILGYWKIKGLVQPTRLLLEYLEEKYEEHLYERDEGDKWRNKKFELGLEFPNLPYYIDGDVKLTQSMAIIRYIADKHNMLGGCPKERAEISMLEGAVLDIRYGVSRIAYSKDFETLKVDFLSKLPEMLKMFEDRLCHKTYLNGDHVTHPDFMLYDALDVVLYMDPMCLDAFPKLVCFKKRIEAIPQIDKYLKSSKYIAWPLQGWQATFGGGDHPPKSDLVPRGSENLYFQGHM**TSKHAHDVCCTSRKEIREQTKHKNNGLILKTGRVNHQSHRPLVVIDDDSSM** | This paper |
| GST-ICD black W22R | GST-thrombin-TEV-ceHPO-30 ICD Y245I W251R F271L | pDK362 | MSPILGYWKIKGLVQPTRLLLEYLEEKYEEHLYERDEGDKWRNKKFELGLEFPNLPYYIDGDVKLTQSMAIIRYIADKHNMLGGCPKERAEISMLEGAVLDIRYGVSRIAYSKDFETLKVDFLSKLPEMLKMFEDRLCHKTYLNGDHVTHPDFMLYDALDVVLYMDPMCLDAFPKLVCFKKRIEAIPQIDKYLKSSKYIAWPLQGWQATFGGGDHPPKSDLVPRGSENLYFQGHM**TSKHAHDVCCTSRKEIREQTKRKNNGLILKTGRVNHQSHRPLVVIDDDSSM** | This paper |
| ICD black W22R | MBP-TEV-ceHPO-30 ICD Y245I W251R F271L, MBP tag removed by TEV cleavage | pDK371 | MKIEEGKLVIWINGDKGYNGLAEVGKKFEKDTGIKVTVEHPDKLEEKFPQVAATGDGPDIIFWAHDRFGGYAQSGLLAEITPDKAFQDKLYPFTWDAVRYNGKLIAYPIAVEALSLIYNKDLLPNPPKTWEEIPALDKELKAKGKSALMFNLQEPYFTWPLIAADGGYAFKYENGKYDIKDVGVDNAGAKAGLTFLVDLIKNKHMNADTDYSIAEAAFNKGETAMTINGPWAWSNIDTSKVNYGVTVLPTFKGQPSKPFVGVLSAGINAASPNKELAKEFLENYLLTDEGLEAVNKDKPLGAVALKSYEEELAKDPRIAATMENAQKGEIMPNIPQMSAFWYAVRTAVINAASGRQTVDEALKDAQTNSSSNNNNNNNNNNLGIEGRISEFENLYFQGHM**TSKHAHDVCCTSRKEIREQTKRKNNGLILKTGRVNHQSHRPLVVIDDDSSM** | This paper |
| ICD-mEGFP | GST-TEV-ceHPO-30 ICD 229-279 – GGS2-mEGFP, GST tag removed by TEV cleavage | pDK184 | MSPILGYWKIKGLVQPTRLLLEYLEEKYEEHLYERDEGDKWRNKKFELGLEFPNLPYYIDGDVKLTQSMAIIRYIADKHNMLGGCPKERAEISMLEGAVLDIRYGVSRIAYSKDFETLKVDFLSKLPEMLKMFEDRLCHKTYLNGDHVTHPDFMLYDALDVVLYMDPMCLDAFPKLVCFKKRIEAIPQIDKYLKSSKYIAWPLQGWQATFGGGDHPPKSDLVPRGSENLYFQGHM**TSKHAHDVCCTSRKEYREQTKWKNNGLILKTGRVNHQSHRPFVVIDDDSSM**GGSGGSMVSKGEELFTGVVPILVELDGDVNGHKFSVSGEGEGDATYGKLTLKFICTTGKLPVPWPTLVTTLTYGVQCFSRYPDHMKQHDFFKSAMPEGYVQERTIFFKDDGNYKTRAEVKFEGDTLVNRIELKGIDFKEDGNILGHKLEYNYNSHNVYIMADKQKNGIKVNFKIRHNIEDGSVQLADHYQQNTPIGDGPVLLPDNHYLSTQSKLSKDPNEKRDHMVLKEKVTAAGITLGMDELYK | This paper |
| ICD | MBP-TEV-ceHPO-30 ICD 229-279, MBP tag removed by TEV cleavage | pDK092 | MKIEEGKLVIWINGDKGYNGLAEVGKKFEKDTGIKVTVEHPDKLEEKFPQVAATGDGPDIIFWAHDRFGGYAQSGLLAEITPDKAFQDKLYPFTWDAVRYNGKLIAYPIAVEALSLIYNKDLLPNPPKTWEEIPALDKELKAKGKSALMFNLQEPYFTWPLIAADGGYAFKYENGKYDIKDVGVDNAGAKAGLTFLVDLIKNKHMNADTDYSIAEAAFNKGETAMTINGPWAWSNIDTSKVNYGVTVLPTFKGQPSKPFVGVLSAGINAASPNKELAKEFLENYLLTDEGLEAVNKDKPLGAVALKSYEEELAKDPRIAATMENAQKGEIMPNIPQMSAFWYAVRTAVINAASGRQTVDEALKDAQTNSSSNNNNNNNNNNLGIEGRISEFENLYFQGHM**TSKHAHDVCCTSRKEYREQTKWKNNGLILKTGRVNHQSHRPFVVIDDDSSM** | This paper |
| DLC8-ICD | DLC8-TEV-ceHPO-30 ICD 229-279-His6 | pDK268 | MSDRKAVIKNADMSEEMQQDAVDCATQALEKYNIEKDIAAYIKKEFDKKYNPTWHCIVGRNFGSYVTHETRHFIYFYLGQVAILLFKSGGSENLYFQGHM**TSKHAHDVCCTSRKEYREQTKWKNNGLILKTGRVNHQSHRPFVVIDDDSSM**HHHHHH | This paper |
| GB1-ICD | GB1-thrombin-ceHPO-30 ICD 229-279-His6 | pDK239 | MQYKLALNGKTLKGETTTEAVDAATAEKVFKQYANDNGVDGEWTYDDATKTFTVTELVPRGS**TSKHAHDVCCTSRKEYREQTKWKNNGLILKTGRVNHQSHRPFVVIDDDSSM**HHHHHH | This paper |
| GB1-homoCC-ICD | GB1-thrombin-homoCC-G-ceHPO-30 ICD 229-279-His6 | pDK336 | MQYKLALNGKTLKGETTTEAVDAATAEKVFKQYANDNGVDGEWTYDDATKTFTVTELVPRGSVKQLEDKVEELLSKNAHLENEVARLKKLVGTSKHAHDVCC**TSRKEYREQTKWKNNGLILKTGRVNHQSHRPFVVIDDDSSM**HHHHHH | This paper (homoCC sequence from O’Shea et al., 1991) |
| sumo-ICD | His10-Sumo-GGS-ceHPO-30 ICD 229-279 | pDK219 | MGHHHHHHHHHHSSGHIEGRHMASMSDSEVNQEAKPEVKPEVKPETHINLKVSDGSSEIFFKIKKTTPLRRLMEAFAKRQGKEMDSLRFLYDGIRIQADQTPEDLDMEDNDIIEAHREQIGGS**TSKHAHDVCCTSRKEYREQTKWKNNGLILKTGRVNHQSHRPFVVIDDDSSM** | This paper |
| GB1-FKBP-ICD | GB1- thrombin-FKBP-GGS3-ceHPO-30 ICD 229-279 – His6 | pDK264 | MQYKLALNGKTLKGETTTEAVDAATAEKVFKQYANDNGVDGEWTYDDATKTFTVTELVPRGSHMVQVETISPGDGRTFPKRGQTCVVHYTGMLEDGKKFDSSRDRNKPFKFMLGKQEVIRGWEEGVAQMSVGQRAKLTISPDYAYGATGHPGIIPPHATLVFDVELLKLEGGSGGSGGSM**TSKHAHDVCCTSRKEYREQTKWKNNGLILKTGRVNHQSHRPFVVIDDDSSM**HHHHHH | This paper |
| GB1-FRB-ICD | GB1- thrombin-FRB-GGS3-ceHPO-30 ICD 229-279 – His6 | pDK265 | MQYKLALNGKTLKGETTTEAVDAATAEKVFKQYANDNGVDGEWTYDDATKTFTVTELVPRGSHMELIRVAILWHEMWHEGLEEASRLYFGERNVKGMFEVLEPLHAMMERGPQTLKETSFNQAYGRDLMEAQEWCRKYMKSGNVKDLTQAWDLYYHVFRRISKQGGSGGSGGSM**TSKHAHDVCCTSRKEYREQTKWKNNGLILKTGRVNHQSHRPFVVIDDDSSM**HHHHHH | This paper |
| GB1-FKBP | GB1- thrombin-FKBP-GGS3-His6 | pDK270 | MQYKLALNGKTLKGETTTEAVDAATAEKVFKQYANDNGVDGEWTYDDATKTFTVTELVPRGSHMVQVETISPGDGRTFPKRGQTCVVHYTGMLEDGKKFDSSRDRNKPFKFMLGKQEVIRGWEEGVAQMSVGQRAKLTISPDYAYGATGHPGIIPPHATLVFDVELLKLEGGSGGSGGSHHHHHH | This paper |
| GB1-FRB | GB1- thrombin-FRB-GGS3-His6 | pDK271 | MQYKLALNGKTLKGETTTEAVDAATAEKVFKQYANDNGVDGEWTYDDATKTFTVTELVPRGSHMELIRVAILWHEMWHEGLEEASRLYFGERNVKGMFEVLEPLHAMMERGPQTLKETSFNQAYGRDLMEAQEWCRKYMKSGNVKDLTQAWDLYYHVFRRISKQGGSGGSGGSHHHHHH | This paper |
| GB1 | GB1-thrombin-His6 | pDK355 | MQYKLALNGKTLKGETTTEAVDAATAEKVFKQYANDNGVDGEWTYDDATKTFTVTELVPRGSLEHHHHHH | This paper |
| GB1-FKBP-ICD ∆5 | GB1- thrombin-FKBP-GGS3-ceHPO-30 ICD Ala 249-253– His6 | pDK274 | MQYKLALNGKTLKGETTTEAVDAATAEKVFKQYANDNGVDGEWTYDDATKTFTVTELVPRGSHMVQVETISPGDGRTFPKRGQTCVVHYTGMLEDGKKFDSSRDRNKPFKFMLGKQEVIRGWEEGVAQMSVGQRAKLTISPDYAYGATGHPGIIPPHATLVFDVELLKLEGGSGGSGGSM**TSKHAHDVCCTSRKEYREQTAAAAAGLILKTGRVNHQSHRPFVVIDDDSSM**HHHHHH | This paper |
| GB1-FRB-ICD ∆5 | GB1-FRB- thrombin-GGS3-ceHPO-30 ICD Ala 249-253– His6 | pDK275 | QYKLALNGKTLKGETTTEAVDAATAEKVFKQYANDNGVDGEWTYDDATKTFTVTELVPRGSHMELIRVAILWHEMWHEGLEEASRLYFGERNVKGMFEVLEPLHAMMERGPQTLKETSFNQAYGRDLMEAQEWCRKYMKSGNVKDLTQAWDLYYHVFRRISKQGGSGGSGGSM**TSKHAHDVCCTSRKEYREQTAAAAAGLILKTGRVNHQSHRPFVVIDDDSSM**HHHHHH | This paper |
| SNAP-FKBP-ICD | SNAP-FKBP-GGS3-ceHPO-30 ICD 229-279-His6 | pDK283 | MDKDCEMKRTTLDSPLGKLELSGCEQGLHEIKLLGKGTSAADAVEVPAPAAVLGGPEPLMQATAWLNAYFHQPEAIEEFPVPALHHPVFQQESFTRQVLWKLLKVVKFGEVISYQQLAALAGNPAATAAVKTALSGNPVPILIPCHRVVSSSGAAGGYEGGLAVKEWLLAHEGHRLGKPGLGPAGIGAPGSMVQVETISPGDGRTFPKRGQTCVVHYTGMLEDGKKFDSSRDRNKPFKFMLGKQEVIRGWEEGVAQMSVGQRAKLTISPDYAYGATGHPGIIPPHATLVFDVELLKLEGGSGGSGGSM**TSKHAHDVCCTSRKEYREQTKWKNNGLILKTGRVNHQSHRPFVVIDDDSSM**HHHHHH | https://www.addgene.org/101137/ and this paper |
| SNAP-FKBP-ICD ∆1 | SNAP-FKBP-GGS3-ceHPO-30 ICD Ala 229-233-His6 | pDK312 | MDKDCEMKRTTLDSPLGKLELSGCEQGLHEIKLLGKGTSAADAVEVPAPAAVLGGPEPLMQATAWLNAYFHQPEAIEEFPVPALHHPVFQQESFTRQVLWKLLKVVKFGEVISYQQLAALAGNPAATAAVKTALSGNPVPILIPCHRVVSSSGAAGGYEGGLAVKEWLLAHEGHRLGKPGLGPAGIGAPGSMVQVETISPGDGRTFPKRGQTCVVHYTGMLEDGKKFDSSRDRNKPFKFMLGKQEVIRGWEEGVAQMSVGQRAKLTISPDYAYGATGHPGIIPPHATLVFDVELLKLEGGSGGSGGSM**AAAAAHDVCCTSRKEYREQTKWKNNGLILKTGRVNHQSHRPFVVIDDDSSM**HHHHHH | https://www.addgene.org/101137/ and this paper |
| SNAP-FKBP-ICD ∆2 | SNAP-FKBP-GGS3-ceHPO-30 ICD Ala 234-238-His6 | pDK313 | MDKDCEMKRTTLDSPLGKLELSGCEQGLHEIKLLGKGTSAADAVEVPAPAAVLGGPEPLMQATAWLNAYFHQPEAIEEFPVPALHHPVFQQESFTRQVLWKLLKVVKFGEVISYQQLAALAGNPAATAAVKTALSGNPVPILIPCHRVVSSSGAAGGYEGGLAVKEWLLAHEGHRLGKPGLGPAGIGAPGSMVQVETISPGDGRTFPKRGQTCVVHYTGMLEDGKKFDSSRDRNKPFKFMLGKQEVIRGWEEGVAQMSVGQRAKLTISPDYAYGATGHPGIIPPHATLVFDVELLKLEGGSGGSGGSM**TSKHAAAAAATSRKEYREQTKWKNNGLILKTGRVNHQSHRPFVVIDDDSSM**HHHHHH | https://www.addgene.org/101137/ and this paper |
| SNAP-FKBP-ICD ∆3 | SNAP-FKBP-GGS3-ceHPO-30 ICD Ala 239-243-His6 | pDK314 | MDKDCEMKRTTLDSPLGKLELSGCEQGLHEIKLLGKGTSAADAVEVPAPAAVLGGPEPLMQATAWLNAYFHQPEAIEEFPVPALHHPVFQQESFTRQVLWKLLKVVKFGEVISYQQLAALAGNPAATAAVKTALSGNPVPILIPCHRVVSSSGAAGGYEGGLAVKEWLLAHEGHRLGKPGLGPAGIGAPGSMVQVETISPGDGRTFPKRGQTCVVHYTGMLEDGKKFDSSRDRNKPFKFMLGKQEVIRGWEEGVAQMSVGQRAKLTISPDYAYGATGHPGIIPPHATLVFDVELLKLEGGSGGSGGSM**TSKHAHDVCCAAAAAYREQTKWKNNGLILKTGRVNHQSHRPFVVIDDDSSM**HHHHHH | https://www.addgene.org/101137/ and this paper |
| SNAP-FKBP-ICD ∆4 | SNAP-FKBP-GGS3-ceHPO-30 ICD Ala 244-248-His6 | pDK315 | MDKDCEMKRTTLDSPLGKLELSGCEQGLHEIKLLGKGTSAADAVEVPAPAAVLGGPEPLMQATAWLNAYFHQPEAIEEFPVPALHHPVFQQESFTRQVLWKLLKVVKFGEVISYQQLAALAGNPAATAAVKTALSGNPVPILIPCHRVVSSSGAAGGYEGGLAVKEWLLAHEGHRLGKPGLGPAGIGAPGSMVQVETISPGDGRTFPKRGQTCVVHYTGMLEDGKKFDSSRDRNKPFKFMLGKQEVIRGWEEGVAQMSVGQRAKLTISPDYAYGATGHPGIIPPHATLVFDVELLKLEGGSGGSGGSM**TSKHAHDVCCTSRKEAAAAAKWKNNGLILKTGRVNHQSHRPFVVIDDDSSM**HHHHHH | https://www.addgene.org/101137/ and this paper |
| SNAP-FKBP-ICD ∆5 | SNAP-FKBP-GGS3-ceHPO-30 ICD Ala 249-253-His6 | pDK316 | MDKDCEMKRTTLDSPLGKLELSGCEQGLHEIKLLGKGTSAADAVEVPAPAAVLGGPEPLMQATAWLNAYFHQPEAIEEFPVPALHHPVFQQESFTRQVLWKLLKVVKFGEVISYQQLAALAGNPAATAAVKTALSGNPVPILIPCHRVVSSSGAAGGYEGGLAVKEWLLAHEGHRLGKPGLGPAGIGAPGSMVQVETISPGDGRTFPKRGQTCVVHYTGMLEDGKKFDSSRDRNKPFKFMLGKQEVIRGWEEGVAQMSVGQRAKLTISPDYAYGATGHPGIIPPHATLVFDVELLKLEGGSGGSGGSM**TSKHAHDVCCTSRKEYREQTAAAAAGLILKTGRVNHQSHRPFVVIDDDSSM**HHHHHH | https://www.addgene.org/101137/ and this paper |
| SNAP-FKBP-ICD ∆6 | SNAP-FKBP-GGS3-ceHPO-30 ICD Ala 254-258-His6 | pDK317 | MDKDCEMKRTTLDSPLGKLELSGCEQGLHEIKLLGKGTSAADAVEVPAPAAVLGGPEPLMQATAWLNAYFHQPEAIEEFPVPALHHPVFQQESFTRQVLWKLLKVVKFGEVISYQQLAALAGNPAATAAVKTALSGNPVPILIPCHRVVSSSGAAGGYEGGLAVKEWLLAHEGHRLGKPGLGPAGIGAPGSMVQVETISPGDGRTFPKRGQTCVVHYTGMLEDGKKFDSSRDRNKPFKFMLGKQEVIRGWEEGVAQMSVGQRAKLTISPDYAYGATGHPGIIPPHATLVFDVELLKLEGGSGGSGGSM**TSKHAHDVCCTSRKEYREQTKWKNNAAAAATGRVNHQSHRPFVVIDDDSSM**HHHHHH | https://www.addgene.org/101137/ and this paper |
| SNAP-FKBP-ICD ∆7 | SNAP-FKBP-GGS3-ceHPO-30 ICD Ala 259-263-His6 | pDK318 | MDKDCEMKRTTLDSPLGKLELSGCEQGLHEIKLLGKGTSAADAVEVPAPAAVLGGPEPLMQATAWLNAYFHQPEAIEEFPVPALHHPVFQQESFTRQVLWKLLKVVKFGEVISYQQLAALAGNPAATAAVKTALSGNPVPILIPCHRVVSSSGAAGGYEGGLAVKEWLLAHEGHRLGKPGLGPAGIGAPGSMVQVETISPGDGRTFPKRGQTCVVHYTGMLEDGKKFDSSRDRNKPFKFMLGKQEVIRGWEEGVAQMSVGQRAKLTISPDYAYGATGHPGIIPPHATLVFDVELLKLEGGSGGSGGSM**TSKHAHDVCCTSRKEYREQTKWKNNGLILKAAAAAHQSHRPFVVIDDDSSM**HHHHHH | https://www.addgene.org/101137/ and this paper |
| SNAP-FKBP-ICD ∆8 | SNAP-FKBP-GGS3-ceHPO-30 ICD Ala 264-268-His6 | pDK319 | MDKDCEMKRTTLDSPLGKLELSGCEQGLHEIKLLGKGTSAADAVEVPAPAAVLGGPEPLMQATAWLNAYFHQPEAIEEFPVPALHHPVFQQESFTRQVLWKLLKVVKFGEVISYQQLAALAGNPAATAAVKTALSGNPVPILIPCHRVVSSSGAAGGYEGGLAVKEWLLAHEGHRLGKPGLGPAGIGAPGSMVQVETISPGDGRTFPKRGQTCVVHYTGMLEDGKKFDSSRDRNKPFKFMLGKQEVIRGWEEGVAQMSVGQRAKLTISPDYAYGATGHPGIIPPHATLVFDVELLKLEGGSGGSGGSM**TSKHAHDVCCTSRKEYREQTKWKNNGLILKTGRVNAAAAAPFVVIDDDSSM**HHHHHH | https://www.addgene.org/101137/ and this paper |
| SNAP-FKBP-ICD ∆9 | SNAP-FKBP-GGS3-ceHPO-30 ICD Ala 269-273-His6 | pDK320 | MDKDCEMKRTTLDSPLGKLELSGCEQGLHEIKLLGKGTSAADAVEVPAPAAVLGGPEPLMQATAWLNAYFHQPEAIEEFPVPALHHPVFQQESFTRQVLWKLLKVVKFGEVISYQQLAALAGNPAATAAVKTALSGNPVPILIPCHRVVSSSGAAGGYEGGLAVKEWLLAHEGHRLGKPGLGPAGIGAPGSMVQVETISPGDGRTFPKRGQTCVVHYTGMLEDGKKFDSSRDRNKPFKFMLGKQEVIRGWEEGVAQMSVGQRAKLTISPDYAYGATGHPGIIPPHATLVFDVELLKLEGGSGGSGGSM**TSKHAHDVCCTSRKEYREQTKWKNNGLILKTGRVNHQSHRAAAAADDDSSM**HHHHHH | https://www.addgene.org/101137/ and this paper |
| SNAP-FKBP-ICD ∆10 | SNAP-FKBP-GGS3-ceHPO-30 ICD Ala 274-279-His6 | pDK321 | MDKDCEMKRTTLDSPLGKLELSGCEQGLHEIKLLGKGTSAADAVEVPAPAAVLGGPEPLMQATAWLNAYFHQPEAIEEFPVPALHHPVFQQESFTRQVLWKLLKVVKFGEVISYQQLAALAGNPAATAAVKTALSGNPVPILIPCHRVVSSSGAAGGYEGGLAVKEWLLAHEGHRLGKPGLGPAGIGAPGSMVQVETISPGDGRTFPKRGQTCVVHYTGMLEDGKKFDSSRDRNKPFKFMLGKQEVIRGWEEGVAQMSVGQRAKLTISPDYAYGATGHPGIIPPHATLVFDVELLKLEGGSGGSGGSM**TSKHAHDVCCTSRKEYREQTKWKNNGLILKTGRVNHQSHRPFVVIAAAAAA**HHHHHH | https://www.addgene.org/101137/ and this paper |
| SNAP-FRB-ICD | SNAP-FRB-GGS3-ceHPO-30 ICD 229-279-His6 | pDK281 | MDKDCEMKRTTLDSPLGKLELSGCEQGLHEIKLLGKGTSAADAVEVPAPAAVLGGPEPLMQATAWLNAYFHQPEAIEEFPVPALHHPVFQQESFTRQVLWKLLKVVKFGEVISYQQLAALAGNPAATAAVKTALSGNPVPILIPCHRVVSSSGAVGGYEGGLAVKEWLLAHEGHRLGKPGLGPAGIGAPGSMELIRVAILWHEMWHEGLEEASRLYFGERNVKGMFEVLEPLHAMMERGPQTLKETSFNQAYGRDLMEAQEWCRKYMKSGNVKDLTQAWDLYYHVFRRISKQGGSGGSGGSM**TSKHAHDVCCTSRKEYREQTKWKNNGLILKTGRVNHQSHRPFVVIDDDSSM**HHHHHH | https://www.addgene.org/101137/ and this paper |
| SNAP-FRB-ICD ∆1 | SNAP-FRB-GGS3-ceHPO-30 ICD Ala 229-233-His6 | pDK302 | MDKDCEMKRTTLDSPLGKLELSGCEQGLHEIKLLGKGTSAADAVEVPAPAAVLGGPEPLMQATAWLNAYFHQPEAIEEFPVPALHHPVFQQESFTRQVLWKLLKVVKFGEVISYQQLAALAGNPAATAAVKTALSGNPVPILIPCHRVVSSSGAVGGYEGGLAVKEWLLAHEGHRLGKPGLGPAGIGAPGSMELIRVAILWHEMWHEGLEEASRLYFGERNVKGMFEVLEPLHAMMERGPQTLKETSFNQAYGRDLMEAQEWCRKYMKSGNVKDLTQAWDLYYHVFRRISKQGGSGGSGGSM**AAAAAHDVCCTSRKEYREQTKWKNNGLILKTGRVNHQSHRPFVVIDDDSSM**HHHHHH | https://www.addgene.org/101137/ and this paper |
| SNAP-FRB-ICD ∆2 | SNAP-FRB-GGS3-ceHPO-30 ICD Ala 234-238-His6 | pDK303 | MDKDCEMKRTTLDSPLGKLELSGCEQGLHEIKLLGKGTSAADAVEVPAPAAVLGGPEPLMQATAWLNAYFHQPEAIEEFPVPALHHPVFQQESFTRQVLWKLLKVVKFGEVISYQQLAALAGNPAATAAVKTALSGNPVPILIPCHRVVSSSGAVGGYEGGLAVKEWLLAHEGHRLGKPGLGPAGIGAPGSMELIRVAILWHEMWHEGLEEASRLYFGERNVKGMFEVLEPLHAMMERGPQTLKETSFNQAYGRDLMEAQEWCRKYMKSGNVKDLTQAWDLYYHVFRRISKQGGSGGSGGSM**TSKHAAAAAATSRKEYREQTKWKNNGLILKTGRVNHQSHRPFVVIDDDSSM**HHHHHH | https://www.addgene.org/101137/ and this paper |
| SNAP-FRB-ICD ∆3 | SNAP-FRB-GGS3-ceHPO-30 ICD Ala 239-243-His6 | pDK304 | MDKDCEMKRTTLDSPLGKLELSGCEQGLHEIKLLGKGTSAADAVEVPAPAAVLGGPEPLMQATAWLNAYFHQPEAIEEFPVPALHHPVFQQESFTRQVLWKLLKVVKFGEVISYQQLAALAGNPAATAAVKTALSGNPVPILIPCHRVVSSSGAVGGYEGGLAVKEWLLAHEGHRLGKPGLGPAGIGAPGSMELIRVAILWHEMWHEGLEEASRLYFGERNVKGMFEVLEPLHAMMERGPQTLKETSFNQAYGRDLMEAQEWCRKYMKSGNVKDLTQAWDLYYHVFRRISKQGGSGGSGGSM**TSKHAHDVCCAAAAAYREQTKWKNNGLILKTGRVNHQSHRPFVVIDDDSSM**HHHHHH | https://www.addgene.org/101137/ and this paper |
| SNAP-FRB-ICD ∆4 | SNAP-FRB-GGS3-ceHPO-30 ICD Ala 244-248-His6 | pDK305 | MDKDCEMKRTTLDSPLGKLELSGCEQGLHEIKLLGKGTSAADAVEVPAPAAVLGGPEPLMQATAWLNAYFHQPEAIEEFPVPALHHPVFQQESFTRQVLWKLLKVVKFGEVISYQQLAALAGNPAATAAVKTALSGNPVPILIPCHRVVSSSGAVGGYEGGLAVKEWLLAHEGHRLGKPGLGPAGIGAPGSMELIRVAILWHEMWHEGLEEASRLYFGERNVKGMFEVLEPLHAMMERGPQTLKETSFNQAYGRDLMEAQEWCRKYMKSGNVKDLTQAWDLYYHVFRRISKQGGSGGSGGSM**TSKHAHDVCCTSRKEAAAAAKWKNNGLILKTGRVNHQSHRPFVVIDDDSSM**HHHHHH | https://www.addgene.org/101137/ and this paper |
| SNAP-FRB-ICD ∆5 | SNAP-FRB-GGS3-ceHPO-30 ICD Ala 249-253-His6 | pDK306 | MDKDCEMKRTTLDSPLGKLELSGCEQGLHEIKLLGKGTSAADAVEVPAPAAVLGGPEPLMQATAWLNAYFHQPEAIEEFPVPALHHPVFQQESFTRQVLWKLLKVVKFGEVISYQQLAALAGNPAATAAVKTALSGNPVPILIPCHRVVSSSGAVGGYEGGLAVKEWLLAHEGHRLGKPGLGPAGIGAPGSMELIRVAILWHEMWHEGLEEASRLYFGERNVKGMFEVLEPLHAMMERGPQTLKETSFNQAYGRDLMEAQEWCRKYMKSGNVKDLTQAWDLYYHVFRRISKQGGSGGSGGSM**TSKHAHDVCCTSRKEYREQTAAAAAGLILKTGRVNHQSHRPFVVIDDDSSM**HHHHHH | https://www.addgene.org/101137/ and this paper |
| SNAP-FRB-ICD ∆6 | SNAP-FRB-GGS3-ceHPO-30 ICD Ala 254-258-His6 | pDK307 | MDKDCEMKRTTLDSPLGKLELSGCEQGLHEIKLLGKGTSAADAVEVPAPAAVLGGPEPLMQATAWLNAYFHQPEAIEEFPVPALHHPVFQQESFTRQVLWKLLKVVKFGEVISYQQLAALAGNPAATAAVKTALSGNPVPILIPCHRVVSSSGAVGGYEGGLAVKEWLLAHEGHRLGKPGLGPAGIGAPGSMELIRVAILWHEMWHEGLEEASRLYFGERNVKGMFEVLEPLHAMMERGPQTLKETSFNQAYGRDLMEAQEWCRKYMKSGNVKDLTQAWDLYYHVFRRISKQGGSGGSGGSM**TSKHAHDVCCTSRKEYREQTKWKNNAAAAATGRVNHQSHRPFVVIDDDSSM**HHHHHH | https://www.addgene.org/101137/ and this paper |
| SNAP-FRB-ICD ∆7 | SNAP-FRB-GGS3-ceHPO-30 ICD Ala 259-263-His6 | pDK308 | MDKDCEMKRTTLDSPLGKLELSGCEQGLHEIKLLGKGTSAADAVEVPAPAAVLGGPEPLMQATAWLNAYFHQPEAIEEFPVPALHHPVFQQESFTRQVLWKLLKVVKFGEVISYQQLAALAGNPAATAAVKTALSGNPVPILIPCHRVVSSSGAVGGYEGGLAVKEWLLAHEGHRLGKPGLGPAGIGAPGSMELIRVAILWHEMWHEGLEEASRLYFGERNVKGMFEVLEPLHAMMERGPQTLKETSFNQAYGRDLMEAQEWCRKYMKSGNVKDLTQAWDLYYHVFRRISKQGGSGGSGGSM**TSKHAHDVCCTSRKEYREQTKWKNNGLILKAAAAAHQSHRPFVVIDDDSSM**HHHHHH | https://www.addgene.org/101137/ and this paper |
| SNAP-FRB-ICD ∆8 | SNAP-FRB-GGS3-ceHPO-30 ICD Ala 264-268-His6 | pDK309 | MDKDCEMKRTTLDSPLGKLELSGCEQGLHEIKLLGKGTSAADAVEVPAPAAVLGGPEPLMQATAWLNAYFHQPEAIEEFPVPALHHPVFQQESFTRQVLWKLLKVVKFGEVISYQQLAALAGNPAATAAVKTALSGNPVPILIPCHRVVSSSGAVGGYEGGLAVKEWLLAHEGHRLGKPGLGPAGIGAPGSMELIRVAILWHEMWHEGLEEASRLYFGERNVKGMFEVLEPLHAMMERGPQTLKETSFNQAYGRDLMEAQEWCRKYMKSGNVKDLTQAWDLYYHVFRRISKQGGSGGSGGSM**TSKHAHDVCCTSRKEYREQTKWKNNGLILKTGRVNAAAAAPFVVIDDDSSM**HHHHHH | https://www.addgene.org/101137/ and this paper |
| SNAP-FRB-ICD ∆9 | SNAP-FRB-GGS3-ceHPO-30 ICD Ala 269-273-His6 | PDK310 | MDKDCEMKRTTLDSPLGKLELSGCEQGLHEIKLLGKGTSAADAVEVPAPAAVLGGPEPLMQATAWLNAYFHQPEAIEEFPVPALHHPVFQQESFTRQVLWKLLKVVKFGEVISYQQLAALAGNPAATAAVKTALSGNPVPILIPCHRVVSSSGAVGGYEGGLAVKEWLLAHEGHRLGKPGLGPAGIGAPGSMELIRVAILWHEMWHEGLEEASRLYFGERNVKGMFEVLEPLHAMMERGPQTLKETSFNQAYGRDLMEAQEWCRKYMKSGNVKDLTQAWDLYYHVFRRISKQGGSGGSGGSM**TSKHAHDVCCTSRKEYREQTKWKNNGLILKTGRVNHQSHRAAAAADDDSSM**HHHHHH | https://www.addgene.org/101137/ and this paper |
| SNAP-FRB-ICD ∆10 | SNAP-FRB-GGS3-ceHPO-30 ICD Ala 274-279-His6 | pDK311 | MDKDCEMKRTTLDSPLGKLELSGCEQGLHEIKLLGKGTSAADAVEVPAPAAVLGGPEPLMQATAWLNAYFHQPEAIEEFPVPALHHPVFQQESFTRQVLWKLLKVVKFGEVISYQQLAALAGNPAATAAVKTALSGNPVPILIPCHRVVSSSGAVGGYEGGLAVKEWLLAHEGHRLGKPGLGPAGIGAPGSMELIRVAILWHEMWHEGLEEASRLYFGERNVKGMFEVLEPLHAMMERGPQTLKETSFNQAYGRDLMEAQEWCRKYMKSGNVKDLTQAWDLYYHVFRRISKQGGSGGSGGSM**TSKHAHDVCCTSRKEYREQTKWKNNGLILKTGRVNHQSHRPFVVIAAAAAA**HHHHHH | https://www.addgene.org/101137/ and this paper |
| SNAP-FKBP | SNAP-FKBP-GGS3-His6 | pDK282 | MDKDCEMKRTTLDSPLGKLELSGCEQGLHEIKLLGKGTSAADAVEVPAPAAVLGGPEPLMQATAWLNAYFHQPEAIEEFPVPALHHPVFQQESFTRQVLWKLLKVVKFGEVISYQQLAALAGNPAATAAVKTALSGNPVPILIPCHRVVSSSGAVGGYEGGLAVKEWLLAHEGHRLGKPGLGPAGIGAPGSMVQVETISPGDGRTFPKRGQTCVVHYTGMLEDGKKFDSSRDRNKPFKFMLGKQEVIRGWEEGVAQMSVGQRAKLTISPDYAYGATGHPGIIPPHATLVFDVELLKLEGGSGGSGGSHHHHHH | https://www.addgene.org/101137/ and this paper |
| SNAP-FRB | SNAP-FRB-GGS3-His6 | pDK280 | MDKDCEMKRTTLDSPLGKLELSGCEQGLHEIKLLGKGTSAADAVEVPAPAAVLGGPEPLMQATAWLNAYFHQPEAIEEFPVPALHHPVFQQESFTRQVLWKLLKVVKFGEVISYQQLAALAGNPAATAAVKTALSGNPVPILIPCHRVVSSSGAVGGYEGGLAVKEWLLAHEGHRLGKPGLGPAGIGAPGSMELIRVAILWHEMWHEGLEEASRLYFGERNVKGMFEVLEPLHAMMERGPQTLKETSFNQAYGRDLMEAQEWCRKYMKSGNVKDLTQAWDLYYHVFRRISKQGGSGGSGGSHHHHHH | https://www.addgene.org/101137/ and this paper |
| SNAP-CapZ | His9-SNAP-CapZ β / CapZ α, from *Gallus gallus* | pDK288 | MGHHHHHHHHHENLYFQGSEFMDKDCEMKRTTLDSPLGKLELSGCEQGLHEIKLLGKGTSAADAVEVPAPAAVLGGPEPLMQATAWLNAYFHQPEAIEEFPVPALHHPVFQQESFTRQVLWKLLKVVKFGEVISYQQLAALAGNPAATAAVKTALSGNPVPILIPCHRVVSSSGAVGGYEGGLAVKEWLLAHEGHRLGKPGLGPAGIGAPGS**MSDQQLDCALDLMRRLPPQQIEKNLSDLIDLVPSLCEDLLSSVDQPLKIARDKVVGKDYLLCDYNRDGDSYRSPWSNKYDPPLEDGAMPSARLRKLEVEANNAFDQYRDLYFEGGVSSVYLWDLDHGFAGVILIKKAGDGSKKIKGCWDSIHVVEVQEKSSGRTAHYKLTSTVMLWLQTNKTGSGTMNLGGSLTRQMEKDETVSDSSPHIANIGRLVEDMENKIRSTLNEIYFGKTKDIVNGLRSIDAIPDNQKYKQLQRELSQVLTQRQIYIQPDN** / **MADFEDRVSDEEKVRIAAKFITHAPPGEFNEVFNDVRLLLNNDNLLREGAAHAFAQYNMDQFTPVKIEGYDDQVLITEHGDLGNGRFLDPRNKISFKFDHLRKEASDPQPEDTESALKQWRDACDSALRAYVKDHYPNGFCTVYGKSIDGQQTIIACIESHQFQPKNFWNGRWRSEWKFTITPPTAQVAAVLKIQVHYYEDGNVQLVSHKDIQDSVQVSSDVQTAKEFIKIIENAENEYQTAISENYQTMSDTTFKALRRQLPVTRTKIDWNKILSYKIGKEMQNA** | https://www.addgene.org/101137/ and this paper |
| Sra1 | His6-TEV-hSra1 (1-1253, FL). His6 tag removed by TEV cleavage | pDK116 | MSYYHHHHHHDYDIPTTENLYFQGA**MAAQVTLEDALSNVDLLEELPLPDQQPCIEPPPSSLLYQPNFNTNFEDRNAFVTGIARYIEQATVHSSMNEMLEEGQEYAVMLYTWRSCSRAIPQVKCNEQPNRVEIYEKTVEVLEPEVTKLMNFMYFQRNAIERFCGEVRRLCHAERRKDFVSEAYLITLGKFINMFAVLDELKNMKCSVKNDHSAYKRAAQFLRKMADPQSIQESQNLSMFLANHNKITQSLQQQLEVISGYEELLADIVNLCVDYYENRMYLTPSEKHMLLKVMGFGLYLMDGSVSNIYKLDAKKRINLSKIDKYFKQLQVVPLFGDMQIELARYIKTSAHYEENKSRWTCTSSGSSPQYNICEQMIQIREDHMRFISELARYSNSEVVTGSGRQEAQKTDAEYRKLFDLALQGLQLLSQWSAHVMEVYSWKLVHPTDKYSNKDCPDSAEEYERATRYNYTSEEKFALVEVIAMIKGLQVLMGRMESVFNHAIRHTVYAALQDFSQVTLREPLRQAIKKKKNVIQSVLQAIRKTVCDWETGHEPFNDPALRGEKDPKSGFDIKVPRRAVGPSSTQLYMVRTMLESLIADKSGSKKTLRSSLEGPTILDIEKFHRESFFYTHLINFSETLQQCCDLSQLWFREFFLELTMGRRIQFPIEMSMPWILTDHILETKEASMMEYVLYSLDLYNDSAHYALTRFNKQFLYDEIEAEVNLCFDQFVYKLADQIFAYYKVMAGSLLLDKRLRSECKNQGATIHLPPSNRYETLLKQRHVQLLGRSIDLNRLITQRVSAAMYKSLELAIGRFESEDLTSIVELDGLLEINRMTHKLLSRYLTLDGFDAMFREANHNVSAPYGRITLHVFWELNYDFLPNYCYNGSTNRFVRTVLPFSQEFQRDKQPNAQPQYLHGSKALNLAYSSIYGSYRNFVGPPHFQVICRLLGYQGIAVVMEELLKVVKSLLQGTILQYVKTLMEVMPKICRLPRHEYGSPGILEFFHHQLKDIVEYAELKTVCFQNLREVGNAILFCLLIEQSLSLEEVCDLLHAAPFQNILPRVHVKEGERLDAKMKRLESKYAPLHLVPLIERLGTPQQIAIAREGDLLTKERLCCGLSMFEVILTRIRSFLDDPIWRGPLPSNGVMHVDECVEFHRLWSAMQFVYCIPVGTHEFTVEQCFGDGLHWAGCMIIVLLGQQRRFAVLDFCYHLLKVQKHDGKDEIIKNVPLKKMVERIRKFQILNDEIITILDKYLKSGDGEGTPVEHVRCFQPPIHQSLASS** | (Ismail et al., 2009) |
| Nap1 | hNap1 (1-1128, FL) | pDK149 | **MSRSVLQPSQQKLAEKLTILNDRGVGMLTRLYNIKKACGDPKAKPSYLIDKNLESAVKFIVRKFPAVETRNNNQQLAQLQKEKSEILKNLALYYFTFVDVMEFKDHVCELLNTIDVCQVFFDITVNFDLTKNYLDLIITYTTLMILLSRIEERKAIIGLYNYAHEMTHGASDREYPRLGQMIVDYENPLKKMMEEFVPHSKSLSDALISLQMVYPRRNLSADQWRNAQLLSLISAPSTMLNPAQSDTMPCEYLSLDAMEKWIIFGFILCHGILNTDATALNLWKLALQSSSCLSLFRDEVFHIHKAAEDLFVNIRGYNKRINDIRECKEAAVSHAGSMHRERRKFLRSALKELATVLSDQPGLLGPKALFVFMALSFARDEIIWLLRHADNMPKKSADDFIDKHIAELIFYMEELRAHVRKYGPVMQRYYVQYLSGFDAVVLNELVQNLSVCPEDESIIMSSFVNTMTSLSVKQVEDGEVFDFRGMRLDWFRLQAYTSVSKASLGLADHRELGKMMNTIIFHTKMVDSLVEMLVETSDLSIFCFYSRAFEKMFQQCLELPSQSRYSIAFPLLCTHFMSCTHELCPEERHHIGDRSLSLCNMFLDEMAKQARNLITDICTEQCTLSDQLLPKHCAKTISQAVNKKSKKQTGKKGEPEREKPGVESMRKNRLVVTNLDKLHTALSELCFSINYVPNMVVWEHTFTPREYLTSHLEIRFTKSIVGMTMYNQATQEIAKPSELLTSVRAYMTVLQSIENYVQIDITRVFNNVLLQQTQHLDSHGEPTITSLYTNWYLETLLRQVSNGHIAYFPAMKAFVNLPTENELTFNAEEYSDISEMRSLSELLGPYGMKFLSESLMWHISSQVAELKKLVVENVDVLTQMRTSFDKPDQMAALFKRLSSVDSVLKRMTIIGVILSFRSLAQEALRDVLSYHIPFLVSSIEDFKDHIPRETDMKVAMNVYELSSAAGLPCEIDPALVVALSSQKSENISPEEEYKIACLLMVFVAVSLPTLASNVMSQYSPAIEGHCNNIHCLAKAINQIAAALFTIHKGSIEDRLKEFLALASSSLLKIGQETDKTTTRNRESVYLLLDMIVQESPFLTMDLLESCFPYVLLRNAYHAVYKQSVTSSA** | (Ismail et al., 2009) |
| WAVE1^∆230WCA^ | MBP-TEV-hWAVE(1) 1-230, MBP tag removed by TEV cleavage | pDK071 | MKIEEGKLVIWINGDKGYNGLAEVGKKFEKDTGIKVTVEHPDKLEEKFPQVAATGDGPDIIFWAHDRFGGYAQSGLLAEITPDKAFQDKLYPFTWDAVRYNGKLIAYPIAVEALSLIYNKDLLPNPPKTWEEIPALDKELKAKGKSALMFNLQEPYFTWPLIAADGGYAFKYENGKYDIKDVGVDNAGAKAGLTFLVDLIKNKHMNADTDYSIAEAAFNKGETAMTINGPWAWSNIDTSKVNYGVTVLPTFKGQPSKPFVGVLSAGINAASPNKELAKEFLENYLLTDEGLEAVNKDKPLGAVALKSYEEELAKDPRIAATMENAQKGEIMPNIPQMSAFWYAVRTAVINAASGRQTVDEALKDAQTNSSSNNNNNNNNNNLGIEGRISEFENLYFQGH**MPLVKRNIDPRHLCHTALPRGIKNELECVTNISLANIIRQLSSLSKYAEDIFGELFNEAHSFSFRVNSLQERVDRLSVSVTQLDPKEEELSLQDITMRKAFRSSTIQDQQLFDRKTLPIPLQETYDVCEQPPPLNILTPYRDDGKEGLKFYTNPSYFFDLWKEKMLQDTEDKRKEKRKQKQKNLDRPHEPEKVPRAPHDRRREWQKLAQGPELAEDDANLLHKHIEVANG** | (Chen et al., 2017) |
| WAVE1^230WCA^ | MBP-TEV-hWAVE(1) 1-230 – GGS6 – VCA, MBP tag removed by TEV cleavage | pDK081 | MKIEEGKLVIWINGDKGYNGLAEVGKKFEKDTGIKVTVEHPDKLEEKFPQVAATGDGPDIIFWAHDRFGGYAQSGLLAEITPDKAFQDKLYPFTWDAVRYNGKLIAYPIAVEALSLIYNKDLLPNPPKTWEEIPALDKELKAKGKSALMFNLQEPYFTWPLIAADGGYAFKYENGKYDIKDVGVDNAGAKAGLTFLVDLIKNKHMNADTDYSIAEAAFNKGETAMTINGPWAWSNIDTSKVNYGVTVLPTFKGQPSKPFVGVLSAGINAASPNKELAKEFLENYLLTDEGLEAVNKDKPLGAVALKSYEEELAKDPRIAATMENAQKGEIMPNIPQMSAFWYAVRTAVINAASGRQTVDEALKDAQTNSSSNNNNNNNNNNLGIEGRISEFENLYFQGH**MPLVKRNIDPRHLCHTALPRGIKNELECVTNISLANIIRQLSSLSKYAEDIFGELFNEAHSFSFRVNSLQERVDRLSVSVTQLDPKEEELSLQDITMRKAFRSSTIQDQQLFDRKTLPIPLQETYDVCEQPPPLNILTPYRDDGKEGLKFYTNPSYFFDLWKEKMLQDTEDKRKEKRKQKQKNLDRPHEPEKVPRAPHDRRREWQKLAQGPELAEDDANLLHKHIEVANGGGSGGSGGSGGSGGSGGSKRHPSTLPVISDARSVLLEAIRKGIQLRKVEEQREQEAKHERIENDVATILSRRIAVEYSDSEDDSEFDEVDWLE** | This paper |
| Abi2 (1-158) | MBP-TEV-hAbi(2) 1-158, MBP tag removed by TEV cleavage | pDK075 | MKIEEGKLVIWINGDKGYNGLAEVGKKFEKDTGIKVTVEHPDKLEEKFPQVAATGDGPDIIFWAHDRFGGYAQSGLLAEITPDKAFQDKLYPFTWDAVRYNGKLIAYPIAVEALSLIYNKDLLPNPPKTWEEIPALDKELKAKGKSALMFNLQEPYFTWPLIAADGGYAFKYENGKYDIKDVGVDNAGAKAGLTFLVDLIKNKHMNADTDYSIAEAAFNKGETAMTINGPWAWSNIDTSKVNYGVTVLPTFKGQPSKPFVGVLSAGINAASPNKELAKEFLENYLLTDEGLEAVNKDKPLGAVALKSYEEELAKDPRIAATMENAQKGEIMPNIPQMSAFWYAVRTAVINAASGRQTVDEALKDAQTNSSSNNNNNNNNNNLGIEGRISEFENLYFQGH**MAELQMLLEEEIPGGRRALFDSYTNLERVADYCENNYIQSADKQRALEETKAYTTQSLASVAYLINTLANNVLQMLDIQASQLRRMESSINHISQTVDIHKEKVARREIGILTTNKNTSRTHKIIAPANLERPVRYIRKPIDYTILDDIGHGVKVSTQ** | (Ismail et al., 2009) |
| Abi2 (1-158)-sortase | MBP-TEV-hAbi(2) 1-158-LPGTG, MBP tag removed by TEV cleavage | pDK255 | MKIEEGKLVIWINGDKGYNGLAEVGKKFEKDTGIKVTVEHPDKLEEKFPQVAATGDGPDIIFWAHDRFGGYAQSGLLAEITPDKAFQDKLYPFTWDAVRYNGKLIAYPIAVEALSLIYNKDLLPNPPKTWEEIPALDKELKAKGKSALMFNLQEPYFTWPLIAADGGYAFKYENGKYDIKDVGVDNAGAKAGLTFLVDLIKNKHMNADTDYSIAEAAFNKGETAMTINGPWAWSNIDTSKVNYGVTVLPTFKGQPSKPFVGVLSAGINAASPNKELAKEFLENYLLTDEGLEAVNKDKPLGAVALKSYEEELAKDPRIAATMENAQKGEIMPNIPQMSAFWYAVRTAVINAASGRQTVDEALKDAQTNSSSNNNNNNNNNNLGIEGRISEFENLYFQGH**MAELQMLLEEEIPGGRRALFDSYTNLERVADYCENNYIQSADKQRALEETKAYTTQSLASVAYLINTLANNVLQMLDIQASQLRRMESSINHISQTVDIHKEKVARREIGILTTNKNTSRTHKIIAPANLERPVRYIRKPIDYTILDDIGHGVKVSTQGGLPGTGG** | This paper |
| (MBP)_2_-Abi2 | 2MBP-Thrombin-StrepII-hAbi(2) 1-158 | pDK119 | MKIEEGKLVIWINGDKGYNGLAEVGKKFEKDTGIKVTVEHPDKLEEKFPQVAATGDGPDIIFWAHDRFGGYAQSGLLAEITPDKAFQDKLYPFTWDAVRYNGKLIAYPIAVEALSLIYNKDLLPNPPKTWEEIPALDKELKAKGKSALMFNLQEPYFTWPLIAADGGYAFKYENGKYDIKDVGVDNAGAKAGLTFLVDLIKNKHMNADTDYSIAEAAFNKGETAMTINGPWAWSNIDTSKVNYGVTVLPTFKGQPSKPFVGVLSAGINAASPNKELAKEFLENYLLTDEGLEAVNKDKPLGAVALKSYEEELAKDPRIAATMENAQKGEIMPNIPQMSAFWYAVRTAVINAASGRQTVDEALKDAQTNSSSLEWSHPQFEKAGGMKIEEGKLVIWINGDKGYNGLAEVGKKFEKDTGIKVTVEHPDKLEEKFPQVAATGDGPDIIFWAHDRFGGYAQSGLLAEITPDKAFQDKLYPFTWDAVRYNGKLIAYPIAVEALSLIYNKDLLPNPPKTWEEIPALDKELKAKGKSALMFNLQEPYFTWPLIAADGGYAFKYENGKYDIKDVGVDNAGAKAGLTFLVDLIKNKHMNADTDYSIAEAAFNKGETAMTINGPWAWSNIDTSKVNYGVTVLPTFKGQPSKPFVGVLSAGINAASPNKELAKEFLENYLLTDEGLEAVNKDKPLGAVALKSYEEELAKDPRIAATMENAQKGEIMPNIPQMSAFWYAVRTAVINAASGRQTVDEALKDAQTNSSSNNNNNNNNNNLGEFLVPRGSWSHPQFEKAGGH**MAELQMLLEEEIPGGRRALFDSYTNLERVADYCENNYIQSADKQRALEETKAYTTQSLASVAYLINTLANNVLQMLDIQASQLRRMESSINHISQTVDIHKEKVARREIGILTTNKNTSRTHKIIAPANLERPVRYIRKPIDYTILDDIGHGVKVSTQ** | This paper |
| HSPC300 | MBP-TEV-hHSPC300, MBP tag removed by TEV cleavage | pDK069 | MKIEEGKLVIWINGDKGYNGLAEVGKKFEKDTGIKVTVEHPDKLEEKFPQVAATGDGPDIIFWAHDRFGGYAQSGLLAEITPDKAFQDKLYPFTWDAVRYNGKLIAYPIAVEALSLIYNKDLLPNPPKTWEEIPALDKELKAKGKSALMFNLQEPYFTWPLIAADGGYAFKYENGKYDIKDVGVDNAGAKAGLTFLVDLIKNKHMNADTDYSIAEAAFNKGETAMTINGPWAWSNIDTSKVNYGVTVLPTFKGQPSKPFVGVLSAGINAASPNKELAKEFLENYLLTDEGLEAVNKDKPLGAVALKSYEEELAKDPRIAATMENAQKGEIMPNIPQMSAFWYAVRTAVINAASGRQTVDEALKDAQTNSSSNNNNNNNNNNLGIEGRISEFENLYFQGH**MGAAMAGQEDPVQREIHQDWANREYIEIITSSIKKIADFLNSFDMSCRSRLATLNEKLTALERRIEYIEARVTKGETLT** | (Ismail et al., 2009) |
| (MBP)_2_-HSPC300 | 2MBP-Thrombin-StrepII-hHSPC300 | pDK118 | MKIEEGKLVIWINGDKGYNGLAEVGKKFEKDTGIKVTVEHPDKLEEKFPQVAATGDGPDIIFWAHDRFGGYAQSGLLAEITPDKAFQDKLYPFTWDAVRYNGKLIAYPIAVEALSLIYNKDLLPNPPKTWEEIPALDKELKAKGKSALMFNLQEPYFTWPLIAADGGYAFKYENGKYDIKDVGVDNAGAKAGLTFLVDLIKNKHMNADTDYSIAEAAFNKGETAMTINGPWAWSNIDTSKVNYGVTVLPTFKGQPSKPFVGVLSAGINAASPNKELAKEFLENYLLTDEGLEAVNKDKPLGAVALKSYEEELAKDPRIAATMENAQKGEIMPNIPQMSAFWYAVRTAVINAASGRQTVDEALKDAQTNSSSLEWSHPQFEKAGGMKIEEGKLVIWINGDKGYNGLAEVGKKFEKDTGIKVTVEHPDKLEEKFPQVAATGDGPDIIFWAHDRFGGYAQSGLLAEITPDKAFQDKLYPFTWDAVRYNGKLIAYPIAVEALSLIYNKDLLPNPPKTWEEIPALDKELKAKGKSALMFNLQEPYFTWPLIAADGGYAFKYENGKYDIKDVGVDNAGAKAGLTFLVDLIKNKHMNADTDYSIAEAAFNKGETAMTINGPWAWSNIDTSKVNYGVTVLPTFKGQPSKPFVGVLSAGINAASPNKELAKEFLENYLLTDEGLEAVNKDKPLGAVALKSYEEELAKDPRIAATMENAQKGEIMPNIPQMSAFWYAVRTAVINAASGRQTVDEALKDAQTNSSSNNNNNNNNNNLGEFLVPRGSWSHPQFEKAGGH**MGAAMAGQEDPVQREIHQDWANREYIEIITSSIKKIADFLNSFDMSCRSRLATLNEKLTALERRIEYIEARVTKGETLT** | This paper |
| GG-(MBP)_2_ | MKI-GGS-TEV-GG-2MBP-TEV, N-terminal and C-terminal regions removed after TEV cleavage | pDK256 | MKIGGSENLYFQG**GGKTEEGKLVIWINGDKGYNGLAEVGKKFEKDTGIKVTVEHPDKLEEKFPQVAATGDGPDIIFWAHDRFGGYAQSGLLAEITPDKAFQDKLYPFTWDAVRYNGKLIAYPIAVEALSLIYNKDLLPNPPKTWEEIPALDKELKAKGKSALMFNLQEPYFTWPLIAADGGYAFKYENGKYDIKDVGVDNAGAKAGLTFLVDLIKNKHMNADTDYSIAEAAFNKGETAMTINGPWAWSNIDTSKVNYGVTVLPTFKGQPSKPFVGVLSAGINAASPNKELAKEFLENYLLTDEGLEAVNKDKPLGAVALKSYEEELAKDPRIAATMENAQKGEIMPNIPQMSAFWYAVRTAVINAASGRQTVDEALKDAQTNSSSNNGGSGGSGGSKTEEGKLVIWINGDKGYNGLAEVGKKFEKDTGIKVTVEHPDKLEEKFPQVAATGDGPDIIFWAHDRFGGYAQSGLLAEITPDKAFQDKLYPFTWDAVRYNGKLIAYPIAVEALSLIYNKDLLPNPPKTWEEIPALDKELKAKGKSALMFNLQEPYFTWPLIAADGGYAFKYENGKYDIKDVGVDNAGAKAGLTFLVDLIKNKHMNADTDYSIAEAAFNKGETAMTINGPWAWSNIDTSKVNYGVTVLPTFKGQPSKPFVGVLSAGINAASPNKELAKEFLENYLLTDEGLEAVNKDKPLGAVALKSYEEELAKDPRIAATMENAQKGEIMPNIPQMSAFWYAVRTAVINAASGRQTVDEALKDAQTNSSSNNNNNNNNNNLGIEGRISEFENLYFQ**GHMLEEFGSSRVDLQASLALAVVLQRRDWENPGVTQLNRLAAHPPFASWRNSEEARTDRPSQQLRSLNGEWQLGCFGG | This paper |
| Sortase 5M | Sortase 5M-His6 | pDK085 | MQAKPQIPKDKSKVAGYIEIPDADIKEPVYPGPATREQLNRGVSFAEENESLDDQNISIAGHTFIDRPNYQFTNLKAAKKGSMVYFKVGNETRKYKMTSIRNVKPTAVEVLDEQKGKDKQLTLITCDDYNEETGVWETRKIFVATEVKLEHHHHHH | <https://www.addgene.org/75144/> |
| Rac1^QP^ | hRac Q61L/P29SΔ4 | pDK077 | **MQAIKCVVVGDGAVGKTCLLISYTTNAFSGEYIPTVFDNYSANVMVDGKPVNLGLWDTAGLEDYDRLRPLSYPQTDVFLICFSLVSPASFENVRAKWYPEVRHHCPNTPIILVGTKLDLRDDKDTIEKLKEKKLTPITYPQGLAMAKEIGAVKYLECSALTQRGLKTVFDEAIRAVLCPPPVKKRKRK** | This paper |
